# Optical Interference for the Guidance of Cryogenic Focused Ion Beam Milling Beyond the Axial Diffraction Limit

**DOI:** 10.1101/2024.11.01.621231

**Authors:** Anthony V. Sica, Magda Zaoralová, Cali Antolini, Daan B. Boltje, Judit J. Penzes, Lilyana M. Malmqvist, Grant Jensen, Jason T. Kaelber, Peter Dahlberg

## Abstract

Using a tri-coincident cryogenic FIB-SEM-LM system, we identify an interferometric optical response that can be used for targeting lamella production to fluorescently labelled structures with accuracy beyond the diffraction limit. We demonstrate the approach using synthetic samples of fluorescent beads embedded in micron-scale droplets of water. We then apply the approach to capture virions inside host cells. Successful targeting is confirmed by cryogenic electron tomography revealing clusters of virions in intracellular vesicles.

## Main Text

Cryogenic electron tomography (Cryo-ET) is a proven approach for uncovering subcellular architecture *in situ*.^1–3^ However, due to the limited penetration of electrons through biological material, the approach is limited to samples far thinner than typical eukaryotic cells.^4^ To amend this, Cryogenic Focused-Ion-Beam (Cryo-FIB) milling uses an ion beam to ablate cellular material and produce sections of a desirable thickness (lamella), typically on the order of 200 nm.^5^ Targeting the milling process via fluorescence has provided a promising route to capture structures of interest. Recently, commercial options for the integration of light microscopy into the Cryo-FIB vacuum chamber have emerged; however, these systems place the light microscope and focused-ion-beam focal positions in different locations within the chamber.^6^ This makes targeting challenging by requiring a registration process to align the fluorescence and Cryo-FIB images, as well as making real-time response impossible. Image registration requires a high density of objects that are observable in both the optical and ion imaging modalities. Typically, large fluorescently labelled polystyrene beads are used for this purpose,^7–9^ but these can obscure labelled structures of interest. Furthermore, the accuracy of guided milling in this manner suffers from registration errors,^10–12^ localization errors due to refractive index mismatch,^13^ and shifts of the sample during milling.^14^ These effects prohibit the routine targeting of rare and small structures to lamella in the axial dimension. An alternative configuration is to match the focal planes of the optical, electron, and ion microscopes. These tri-coincident systems, such as the ENZEL used here,^15^ can monitor fluorescence in real-time while milling (Figure S2). This ability removes the axial registration problem. So long as the object of interest is significantly larger in axial extent than the target lamella thickness, the object can be captured by halting milling from one direction as fluorescence is observed to dim.^16^ However, this is partially destructive and does not permit the capture of objects on the order of the lamella thickness or smaller. Here, we will show how this limitation can be overcome using an additional interferometric signal to direct milling with no loss of the target structure.

We begin with an artificial system for observation of optical effects during milling. Using a modified plunge freezer, fluorescent polystyrene beads embedded in micron scale water droplets were deposited on the surface of EM grids, Figure 1a and Methods. Frozen droplets were then milled in cleaning cross-section mode (i.e. a line of ions was scanned repeatedly, and that line was slowly advanced axially through the sample). The approach of the ion beam could travel from the bottom of the grid up, termed “bottom-up” milling, or from the top of the droplet down, termed “top-down” milling, Figure 1b. In top-down milling, prominent oscillations in fluorescent intensity are observed as the top surface approaches a fluorescent target. These oscillations grow in amplitude until a rapid drop is seen due to milling through the target, Figure 1d. Oscillations are due to interference between the incoming excitation and its Fresnel reflection off the interface at the top of the sample, see Figure S2 and SI.^17^ Critically, these oscillations inform on the distance between the fluorescent object of interest and the milling surface.

**Figure 1:**
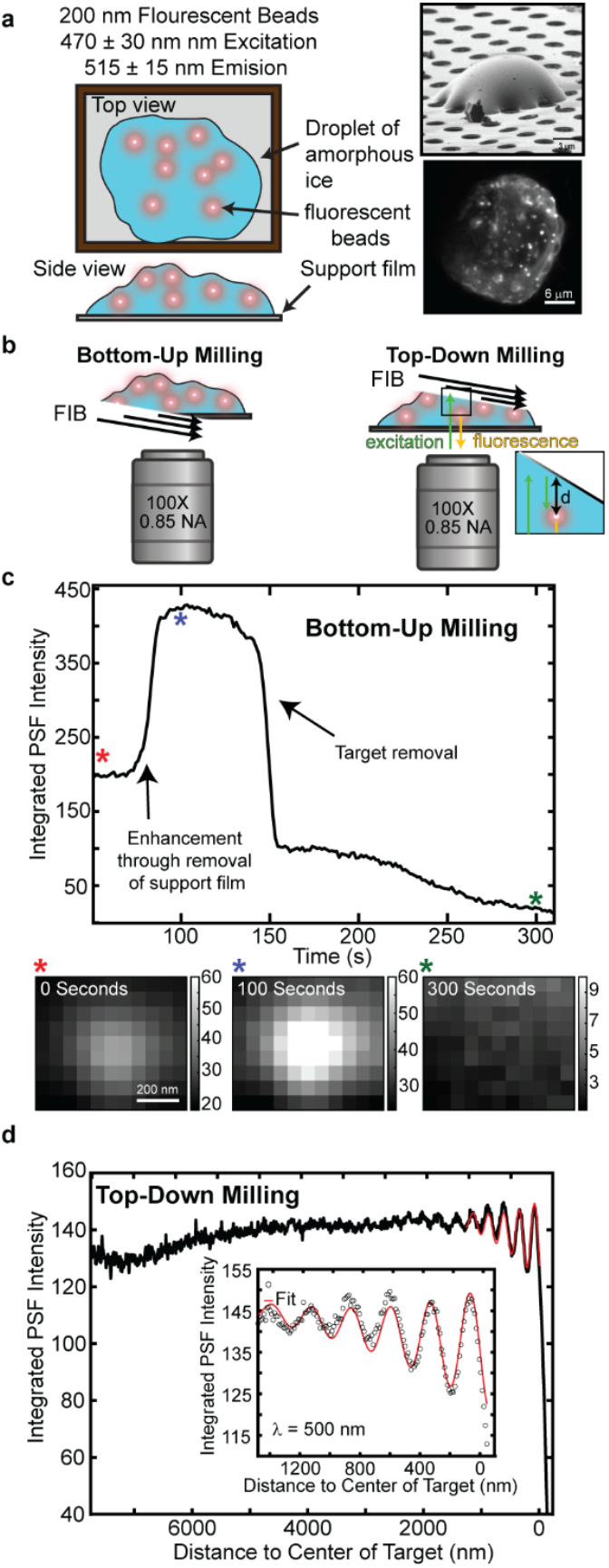
*In vitro* test system for developing fluorescence based interferometric FIB guidance. a.) Fluorescent beads embedded in droplets were sprayed and frozen on a grid. Cartoon depiction, along with an ion and fluorescent image of one droplet. b.) Cartoon depiction of either bottom-up or top-down milling that can be performed while monitoring fluorescence. c.) When milled bottom-up an enhancement in the fluorescence can be seen as the absorptive support film is removed.^31^ Corresponding ROI’s can be seen above to show this enhancement at different times indicated by the colored asterixis. d.) Top-down milling resulted in oscillations in brightness as the surface of the lamella approaches the embedded fluorescent beads. The oscillations were then fit to a model function shown in the inset and are consistent with the excitation wavelength.

To exploit this interference phenomena to direct milling, the fluorescent point-spread-function (PSF) of a sub-diffraction-limited structure of interest is fit to a two-dimensional symmetric Gaussian in real-time. Fitting helps isolate target fluorescence from background fluorescence and more clearly exhibits oscillations as a function of the distance between the interface and the object of interest. Assuming the object originates from a point source, the intensity of oscillations, *I*, as a function of position, *d*, can be fit to the following function:

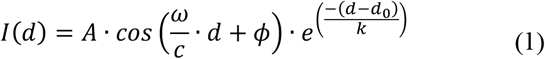

where *A* is the peak amplitude of the oscillations, *ϕ* is the phase, *d*_0_ is an offset fit parameter, *k* is the rate of enhancement in fringe contrast related to the coherence length of the LED-based excitation source, *c* is the speed of light, and *ω* is the frequency of the oscillation. See the SI text for a derivation of equation 1. The recovered frequency in equation 1 is corrected to the optical frequency in vacuum through the following equation

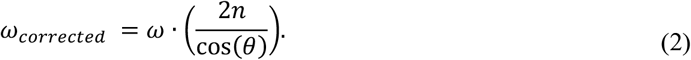

Where n is the index of refraction of amorphous ice^18^ and *θ* is angle of incidence of the FIB relative to the optical axis of the light microscope, with 90 degrees being parallel to the optical axis. In our system this angle is ∼10 ± 2 degrees.

To guide the milling process using the interferogram, it is critical to know the expected amplitude and phase of the oscillation as we approach the target. These factors can be determined from the index of refraction of the two materials at the interface and the angle of incidence as shown by the Fresnel equations^17^, see SI. The dominant source of the reflection could be an interface between amorphous ice and vacuum, or the interface between the amorphous ice and the gallium-doped damage layer at the top surface of the lamella. Based on the amplitudes and phases of the observed interference pattern, we attribute the effect largely to the interface with the damage layer, see SI. Because the amplitude of the reflected wave depends on the damage layer, it varies from sample to sample as a function of the milling parameters such as voltage. Therefore, to guide the milling process with this effect, we first mill through a sacrificial target of interest to record the amplitude prior to milling a proper lamella. Then, using the recovered amplitude and the same milling parameters, subsequent targets could be milled with the oscillations fit to equation 1 and the appropriate trough at which to halt milling could be determined in real-time before milling the target of interest. This information is relayed to the user by a custom Python-based GUI running on the acquisition computer, Figure S4.

We validated this technique with biological samples by targeting adeno-associated virus (AAV) infecting human cells.^19^ AAV is a parvovirus with icosahedral capsids 26 nm in diameter that is most notable for its use in human gene therapy.^20^ Like all parvoviruses, it is internalized by the human cell in endosomes but eventually escapes membrane-bound organelles in a process that (for AAV) involves the phospholipase activity of the sometimes-sequestered N-terminus of capsid protein VP1.^21^ Researchers have postulated that Cryo-ET based observation of AAV as it moves through, and out of, the endomembrane system will unlock mechanistic questions that cannot be reliably recapitulated *in vitro*.^22^ Unfortunately, in contrast to the process of viral assembly, wherein virions are abundant in the infected cell and optical targeting is not typically required^23^, viral trafficking and genome delivery processes necessitate precise ion beam targeting because the small virion in the large mammalian cell constitutes a “needle in a haystack.”

As we had done for fluorescent beads, we first performed top-down milling through several fluorescently labelled AAV-transduced samples (see Methods) to determine the amplitude of oscillations before the virions were ablated. Then, using this value, we directed milling to virion clusters in the main cell body. Tomography of optically-targeted lamellae in HeLa cells at 8 hours post-infection (h.p.i.) revealed clusters of virus, seen predominantly in membranous vesicles; some of these vesicles had a dozen or more concentric bilayers, indicating a lysosomal subtype. Figure 2 and supplementary media show a representative example of interferometrically guided FIB milling. The resulting tomogram shows a densely packed, membrane-bound cluster of virions centered ∼80 nm from the top lamella surface.

**Figure 2:**
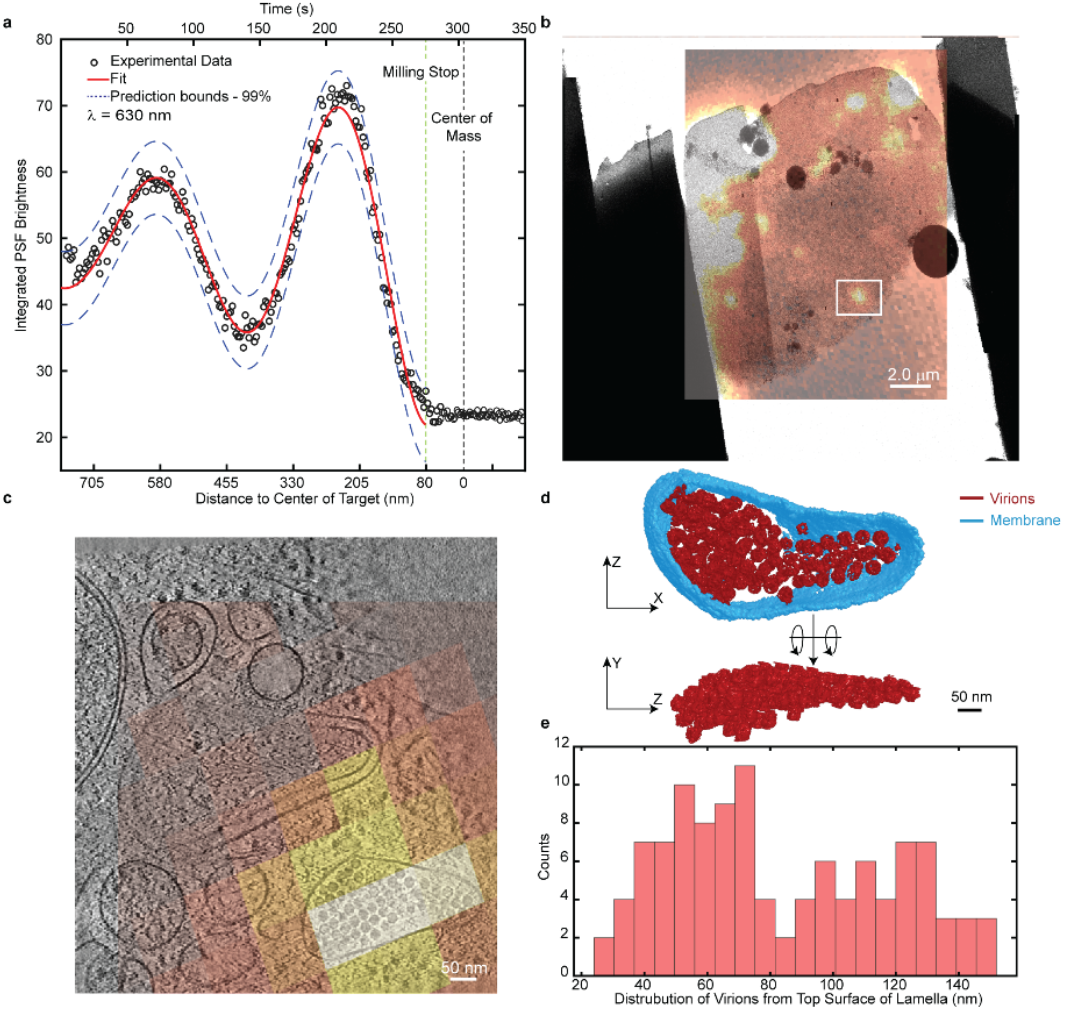
Demonstration of targeted milling to AAV virions. a.) Integrated PSF brightness of target fluorescent puncta while milling. The oscillations are fit using a model function which predicts the stop point for milling. b.) Montage EM showing entire lamella with the corresponding fluorescence overlaid. Region where tomography was measured is shown with the boxed ROI in white. c.) High magnification overlay of the final fluorescent image and a slice from the tomographic reconstruction. d.) Annotation of virions and encapsulating membrane from the tomogram in (c). e.) Histogram of virions in (d) as a function of distance from the top of the lamella with an average distance of ∼80 nm.

Here, we have demonstrated the ability to use fluorescence-based interferometric effects to guide cryoFIB milling of biological systems far beyond the axial diffraction limit. The interferometric approach requires no external fiducials and minimal registration of fluorescence and FIB images. Instead, it uses real-time fluorescence monitoring of a structure of interest to direct the milling process beyond the diffraction limit of light. So long as an object can be distinguished in wide-field fluorescence and the object is not within several hundred nanometers of the top surface of the frozen sample, then interferometric guidance can be performed to direct the milling (see SI for a more detailed discussion of limitations). This approach opens new possibilities to observe small and rare structures previously out of reach for Cryo-FIB milling and Cryo-ET approaches.

## Acknowledgements

The authors would like to thank Chensong Zhang and Lydia-Marie Joubert from the SCSC NIH tomography center for support throughout the project, Jessica Shivas for technical support, and Ash Sueh Hua for clarifying edits. Funding was provided in part by REGENXBIO Inc. (to J.T.K.), and NIGMS grant no. R21GM140345 (J.T.K.), NIAID grant no. RO1AI127401 (GJJ), Department of Energy, Office of Science, Office of Biological and Environmental Research, under Contract No. DE-AC02-76SF00515 FWP 100883 (PDD), and grant 2021-234593 from the Chan Zuckerberg Initiative DAF, an advised fund of Silicon Valley Community Foundation.

## Methods

### Sample preparation

#### Spray frozen fluorescent bead samples

A gas dynamic virtual nozzle (GDVN) previously described in Yoniles et al.^24^ was used to aerosolize a 0.1% solids aqueous sample of 200 nm yellow-green (505/515 nm) polystyrene beads (FluoSpheres™ F8811). Briefly, the GDVN was placed approximately 7 cm perpendicularly to the grid’s path of plunging allow for the aerosolized sample to be sprayed onto the grid before freezing. A total flow rate of 0.14 ml/min and intentionally non-glow discharged copper Quantifoil grids (EMS Q2100CR2) were used to produce 10 – 20 µm droplets with high contact angles. The same was done to aerosolize the large stoke-shift 0.002% solids aqueous sample of 40 nm yellow-red (488/645 nm) Streptavidin-Labeled Microspheres (TransFluoSpheres™ T10711), but at lower percent solids to keep the density of fluorescent objects nearly constant between samples.

#### AAV infected host cells

AAV2 non-replicating virions containing the GFP gene in lieu of *rep/cap*^19^ were labelled in 1× borate buffer (1 M, pH 8.5) in a 15:1 dye to protein ratio with either Alexa Fluor 488 (ThermoFisher) or Janelia Fluor 669 (Tocris) covalently linked NHS ester dye. Stochastic lysine labeling of parvoviruses has minimal impact on infectivity at low multiplicities of labeling but affects the virus as the number of dyes per capsid increases. Preliminary room-temperature confocal microscopy, using Alexa Fluor 488 NHS ester labeled non-replicating AAV2 virions, indicated that the brightness of the dye and wavelength difference from the cellular autofluorescence peak were necessary to detect single virions inside cells. The high autofluorescence detected in the ∼500 nm emission spectrum warranted the switch to the fluorescent dye in the far-red spectrum for the cryogenic fluorescent experiments.

The labeled virions were re-purified on a sucrose gradient in 1× TNTM pH 8 (50 mM Tris pH 8, 100 mM NaCl, 0.2% Triton X-100, 2 mM MgCl_2_) and dialyzed into 1× PBS. For confocal microscopy 0.3×10^6^ HeLa cells were seeded on a glass coverslip and incubated overnight. The culture was pre-chilled for 20 min at 4°C prior to being infected with fluorescently labeled AAV2 at the MOI of 10^4^. Following 30 min of incubation at 4°C to allow the virus particles to attach without being endocytosed, the culture was placed back into the incubator for 20 hours at 37°C with 5% CO_2_. Infected cells were stained with Hoechst stain in 1:2000 concentration and fixed in 2% paraformaldehyde for 30 min. The coverslips were mounted and imaged by a Leica SP8 confocal microscope (Leica Microsystems), Figure S5. For cryogenic imaging, glow discharged Quantifoil gold grids with a holey SiO_2_ support film were disinfected in 70% isopropanol. Following their placement into a 35 mm culturing dish and three washes in 1× PBS, 0.5×10^6^ HeLa or HEK293A cells were seeded onto the disinfected grids and incubated overnight. The cells were pre-chilled and infected with JF-669 labeled AAV2 at the MOI of 10^5^ and incubated as above. Following Hoechst staining, the grids were back-blotted and plunge frozen by a Leica EM GP plunge freezer in liquid ethane.

### Tri-coincident FIB Milling System

The focus ion beam (FIB) milling system used for all studies was the Enzel system initially developed by Delmic installed on an Aquilos 2 FIB-SEM system from ThermoFisher Scientific. Within the Enzel system the stage remains in the same imaging plane while maintaining the ability to simultaneously take FIB, SEM, and light microscopy data. A recreated cartoon depiction of this is seen in Figure S2a along with an image of the inside of the Enzel (Figure S2b). The FIB is equipped with a gallium ion source. Additionally, a gas injection system (GIS) of platinum was layered on all samples prior to milling. More information on the optics employed can be found in a previous publication.^15^

### Software Interface with Enzel System

The Enzel System from Delmic has a tri-coincident geometry, allowing for simultaneous fluorescence and FIB to be performed. To integrate these operations, a Python-based GUI was developed to interface with existing Delmic software. A sample view can be seen in Figure S4. Common controls such as commands to the camera, recording images, LED and filter control, and objective stage movement are integrated into the software. Newly added is the ability to define region of interests (ROI),monitor the average fluorescence, and—if desired—fit the PSF to a 2D Gaussian and integrate and record that fitting. Initial parameters are adjusted to optimize fitting routines and images are continuously saved. As milling progresses, the software will actively fit intensity traces to the model function and extrapolate to when a desired contrast and phase will be reached. Once that condition is reached, the software indicates it is time to stop milling to capture the target. The milling is then stopped manually through simultaneously operating the ion beam using the xT UI interface (ThermoFisher Scientific). The lamella is then cleaned to reach an optimal thickness of ∼200 nm by bottom-up milling.

### Cryogenic Electron Tomography

Tomograms were collected using a 300 keV transmission electron microscope (G2 Titan Krios™) with images recorded using a direct electron detection camera Falcon4i with a Selectris Imaging filter (Thermo Fisher Scientific) and a 10 eV slit width. Images were taken with an exposure time of 2.5 seconds corresponding to 3.5e^−^/Å^2^ dose per projection and a pixel size of 2.5 Å. The integrated images were saved in the mrc format, and individual frames were saved in the eer format.^25^ The tilt series were taken with a dose-symmetric scheme,^26^ starting at a -10 degrees to compensate for the physical pre-tilt of the lamella, imaging with a range of -60 to 40 degrees incremented at 2.5-degree steps. The resulting total dose was approximately 144 e^−^/Å^2^. Reconstruction was done using IMOD and the resulting image binned by 4 reducing pixel size to 9.8 Å.

The eer frames were converted to a tif format (Relion command - convert to tif)^27^, corrected for local and global motion (MotionCor2)^28^ using an option to split even and odd frames that were subsequently used for denoising (CryoCare)^29^. The aligned tilt series were reconstructed to a volume (IMOD)^30^, denoised with CryoCare, and annotated using Dragonfly.

### Image Registration

Fluorescence microscopy was registered to the Cryo-ET data using a series of affine transformations like methods described previously.^31–33^ First the optical images were registered to a low magnification EM image using any features identifiable in both imaging modalities, such as ice or defects in the lamella. These control point pairs were used to compute a projective transformation carrying the optical data into the low-magnification EM space. This low-magnification EM space was then registered with the Cryo-ET space by identifying unique features, such as subcellular structures and ice contamination, visible in both the low-magnification image and the z-projection of the Cryo-ET reconstructions. From these control point pairs a similar transformation was computed to carry the low-magnification EM micrograph to the Cryo-ET space. Application of the projective and similar transformations sequentially to the optical data carries the fluorescence micrographs into the Cryo-ET space.

