## Supplemental Information for "Optical Interference for the Guidance of Cryogenic Focused Ion Beam Milling Beyond the Axial Diffraction Limit"

**Supplementary Text:**

**Interference is a function of excitation wavelength not emission**

Polystyrene fluorescent beads of 200 nm and 40 nm diameter respectively doped with either small Stokes shift (ThermoFisher F8811) or large Stokes shift (ThermoFisher T10711) dyes were milled through in the top-down geometry. The sample with large Stokes shift dye provides a large difference between excitation and emission wavelengths and makes assignment of the interference to the excitation source obvious. Figure S1a shows a large sampling of fluorescent beads where the fluorescence intensity was monitored with a constant top-down milling rate. The oscillations in the traces were then aligned and averaged. The resulting average was then fit to the model function described in the main text (equation 1). The small Stokes shift beads were excited with 470 ± 10 nm and emission collected at 515 ± 15 nm. This yielded a fit with a corresponding periodicity of 500 ± 3.73 nm. Similarly, the process was then repeated for smaller 40 nm beads with the large Stokes shift dye. The dye has a peak excitation of 490 nm and its emission peaks at 645 nm. The large Stokes shift beads were excited with 470 ± 10 nm and emission collected at 698 ± 45 nm. Again, the averaged trace was fit to the same function and a wavelength of 502 ± 7.50 nm was measured (Figure S1b). From this result, we attribute the source of the interference to the excitation light. Interestingly, in the traces of the 40 nm beads, there can be seen a deterioration in the last oscillation. We attribute this to damage from the milling process as the surface of the lamella approaches the fluorescent object.

### Interferometric signal from Fresnel-like reflection

To understand the source of the interferometric signal, we turn to the Fresnel equation for reflection. Our relatively low NA objective leads to much of the light being P polarized at the interface. Thus, we can describe the reflection coefficient, which is the fraction of reflected incident light intensity, as[1–5]

$$R = \left|\frac{n_2 \cos\theta_i - n_1 \cos\theta_t}{n_2 \cos\theta_i + n_1 \cos\theta_t}\right|^2. \tag{1}$$

where $\theta_i$ is the angle of incidence of the incoming light, $\theta_t$ is the angle of refraction of the transmitted light, and $n_1$ is the index of refraction of the amorphous ice and $n_2$ is the refractive index of material at the interface, either vacuum or the gallium doped damage layer, see Figure S2. Snell's law gives the relation between incident angle and the angle of refraction as

$$n_1 \sin(\theta_i) = n_2 \sin(\theta_t). \tag{2}$$

which can be rearranged to determine $\theta_t$ as

$$\operatorname{asin}\left(\frac{n_1}{n_2}\sin(\theta_i)\right) = \theta_t. \tag{3}$$

Substituting equation 3 into equation 1 yields

$$R = \left|\frac{n_2 \cos\theta_i - n_1 \cos\left(\operatorname{asin}\left(\frac{n_1}{n_2}\sin(\theta_i)\right)\right)}{n_2 \cos\theta_i + n_1 \cos\left(\operatorname{asin}\left(\frac{n_1}{n_2}\sin(\theta_i)\right)\right)}\right|^2. \tag{4}$$

Here, the incident angle is well known from the geometry of the setup, 10° ± 2°; the index of refraction of amorphous ice is ~1.26 at 550 nm[6] with the key unknown variable in equation 4 being $n_2$. We will return to equation 4 later, when computing expected amplitudes of the interferogram. From here, we turn to interferometry where the field at a position *d* and time *t* below the lamella is given as[2,7]

$$E_d(t) = E_i(t) + E_r(t + \tau + \phi). \qquad (5)$$

Here, $\tau$ is the time delay due to the round-trip travel from the fluorescent object and back, while $\phi$ is an additional phase factor induced by reflection. If $n_1 < n_2$ then $\phi$ gives a phase delay of π; otherwise $\phi = 0$. The intensity of this field is given as

$$I_d(t) = \frac{1}{2}\epsilon_o c\, |E_d(t)|^2 = \frac{1}{2}\epsilon_o c |E_i(t) + E_r(t + \tau + \phi)|^2. \qquad (6)$$

which expanded gives

$$I_d(t) = \frac{1}{2}\epsilon_o c \left[|E_i(t)|^2 + |E_r(t + \tau + \phi)|^2 + 2Re\big(E_i^*(t)E_r(t + \tau + \phi)\big)\right] \qquad (7)$$

where c is the speed of light and $\epsilon_o$ is the permittivity in a vacuum. The time-average of the two leading homodyne components of equation 7 lead to a constant intensity as a function of *d*, but the cross-term results in oscillations in intensity as the fields $E_i\, E_r$ constructively and destructively interfere as a function of *d*. This oscillatory component is

$$\Delta I_d(t) = \epsilon_o c\, Re\big(E_i^*(t)E_r(t + \tau + \phi)\big). \qquad (8)$$

This can be written as

$$\Delta I_d(t) \propto \sin\left(\frac{2\pi ct}{\lambda}\right) \sin\left(\frac{2\pi ct}{\lambda} + \frac{2d}{\lambda} + \Phi\right) \qquad (9)$$

where $\lambda$ is the excitation wavelength and Φ is either $\pi$ or 0 depending on the relative indices of refraction. Using a product to sum identity, equation 9 can be expanded as:

$$\Delta I_d(t) \propto \frac{1}{2}\left[cos\left(-\left(\frac{2d}{\lambda}+\Phi\right)\right) - cos\left(\left(\frac{4\pi ct}{\lambda}+\frac{2d}{\lambda}+\Phi\right)\right)\right]. \quad (10)$$

Considering the time-averaged field, the second term goes to zero and can be further simplified to:

$$\overline{\Delta I_d} \propto \frac{1}{2}\left[cos\left(\frac{2d}{\lambda}+\Phi\right)\right]. \quad (11)$$

Equational 11 shows that the intensity of the field at position *d* below the lamella surface will oscillate as a function of *d.* However, this assumes an infinitely coherent source. To account for the limited spatial and temporal coherence of the LED-based excitation source, we have an additional exponential to artificially dampen the signal where *k* is the rate of this dampening. This value can be rigorously calculated from the spectral characteristics of the light or taking the autocorrelation of the cross-term in equation 11.[3,4,8]

$$\overline{\Delta I_d} \propto \frac{1}{2}\left[cos\left(\frac{2d}{\lambda}+\Phi\right)\right] * e^{\frac{-d}{k}}. \quad (12)$$

Equation 12 directly relates to equation 1 in the main text.

The interferograms expected from equation 12 are simulated in Figure S3. In this simulation, the fluorescent object was a uniformly fluorescent sphere of 40 nm diameter. The phase factor $\Phi$ is 0 when we consider an interface of amorphous ice and vacuum (Figure S3a), creating a positive amplitude and ending on a peak. Conversely, a phase of $\pi$ is added when assuming the dominating interface to be the gallium doped damaged layer and amorphous ice (Figure S3b), creating a negative amplitude and ending on a trough. Real data from a sample trace of milling through fluorescently labelled virions (Figure 2 from the main text) is overlayed on

Figure S3b to demonstrate the reflections being dominated by this gallium-ice interface. With contamination of gallium seen as high as ~30%, it is not unreasonable to attribute the reflection to gallium deposition in the top layer causing a low to high index reflection.[9] Alongside this one could also look at the fringe contrast seen in the interferences. Using the index of refraction of amorphous ice and vacuum in equation 4, one would expect a reflection coefficient of ~0.013, which gives fringe contrast ~11%. With the contrasts varying from sample to sample and being as high as 34% it becomes more likely that reflections are dominated by gallium contamination rather than an amorphous ice to vacuum interface. An interesting prospect would be to see how these trends might change with the use of different ion sources such as a plasma-based FIB system.

**Necessary criteria for interferometrically guided milling**

To use this approach, two criteria must be met. First, the fluorescent object of interest cannot be within a half wavelengths distance to the top surface of the lamella, in reference to the excitation wavelength. Second, the object must be distinguishable using widefield fluorescence microscopy. The first criterion is straightforward: the object must be far enough from the surface so that top-down milling can estimate the amplitude of oscillation. Without this, it is impossible to determine when to stop milling. The second criterion is difficult to quantify in terms of brightness and copy number of the fluorescent label. In widefield microscopy, especially of thick samples like a mammalian cell, the ability to distinguish a fluorescent puncta from background will depend on both the levels of background from autofluorescence and out-of-focus fluorescence as well as the brightness of the labelled structure. With low enough backgrounds, even single molecules can be observed. Under cryogenic conditions single molecules have been demonstrated to be photostable for tens of minutes, potentially long enough to direct the production of a lamella.[10] While this background is difficult to achieve, it is not inconceivable for certain samples in certain wavelength regimes. As a guiding principle, if the object can be seen in a stand-alone cryogenic light microscope in widefield, then this approach should be able to direct lamella production to preserve the structure for subsequent cryoET.

### Supplementary Figures:

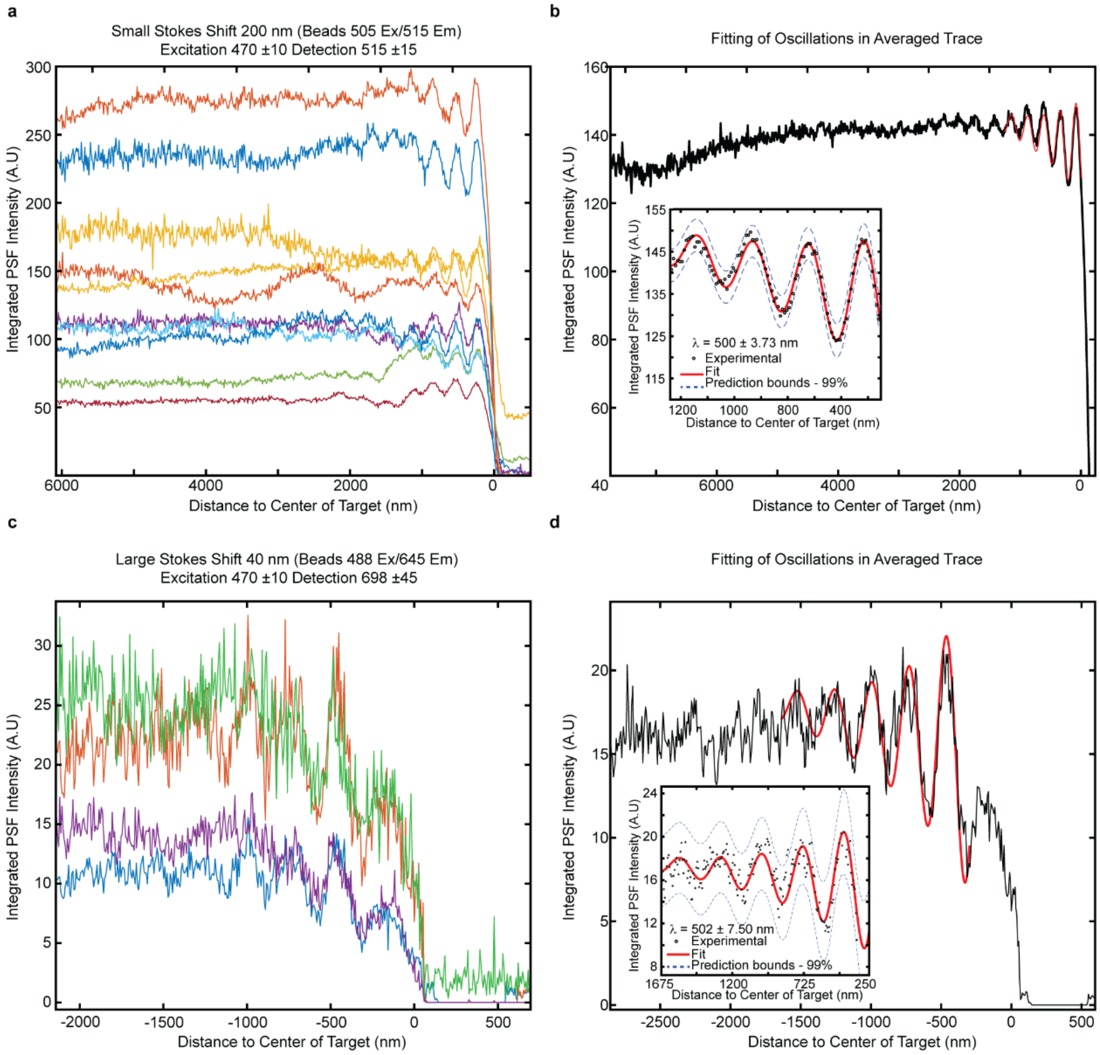


Figure S1: a.) The integrated fluorescence intensity from 200 nm fluorescence beads (470 nm excitation/514 nm emission) sampled across a droplet exhibited consistent oscillations during top-down milling. Each trace is from a separate individual bead. Traces have been aligned by the position of the beads determined by the loss of fluorescence to show the consistency of oscillations. The time axis is relative to each bead. b.) All fluorescent traces were aligned and averaged. The oscillations were fit to a model function to determine the periodicity of interferogram to be 500 ± 3.73 nm. c.) Similarly, 40 nm beads with a larger Stokes shift (480 nm excitation/645 nm emission) were also sampled with consistent oscillations. d.) The traces were aligned, averaged, and fit to the same model function. The resulting periodicity of 502 ± 7.50 nm attributes the source of the interference to be the incoming excitation light and not the fluorescence.

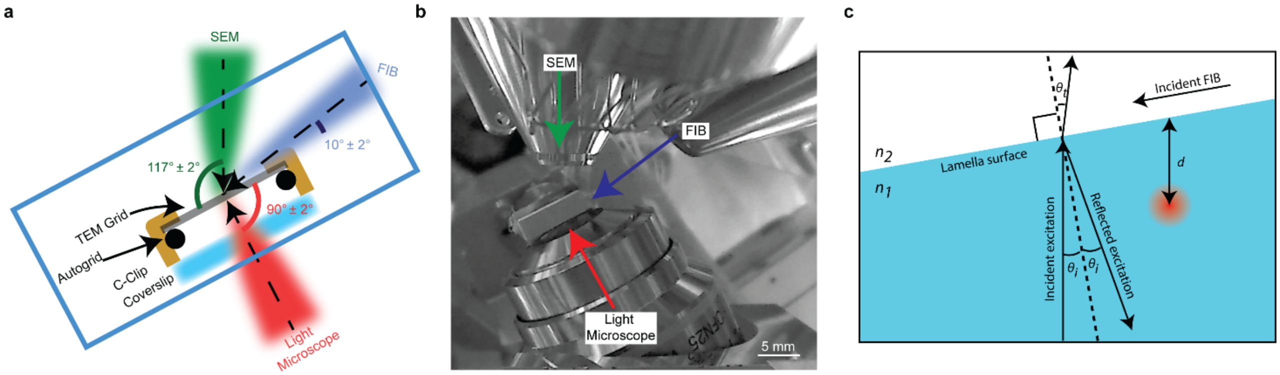


Figure S2: a.) Cartoon depiction of the tri-coincident Enzel system, b.) along with an image of inside the vacuum chamber. Figures 1a and b recreated from ref[4]. c.) Diagram of interface of reflection showing the necessary geometry to recreate interferometric signal

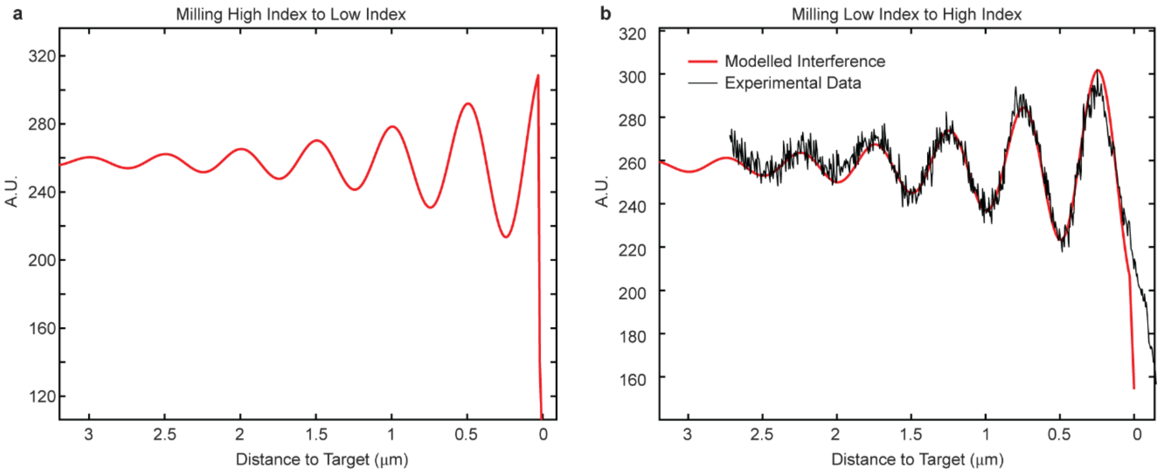


Figure S3: a.) Simulated interference with high to low index interface creating a reflection field that is in phase with the incoming field resulting in constructive interference prior to milling through the object. b.) Interferometric signal created by a low to high index interface producing a 180 degree phase change in the reflected field relative to the incoming field, resulting in destructive interference prior to milling through the object. A fluorescence intensity trace from milling through virions is shown fit to the simulated data to emphasize the likeness that the low to high index signal is to experimental data.

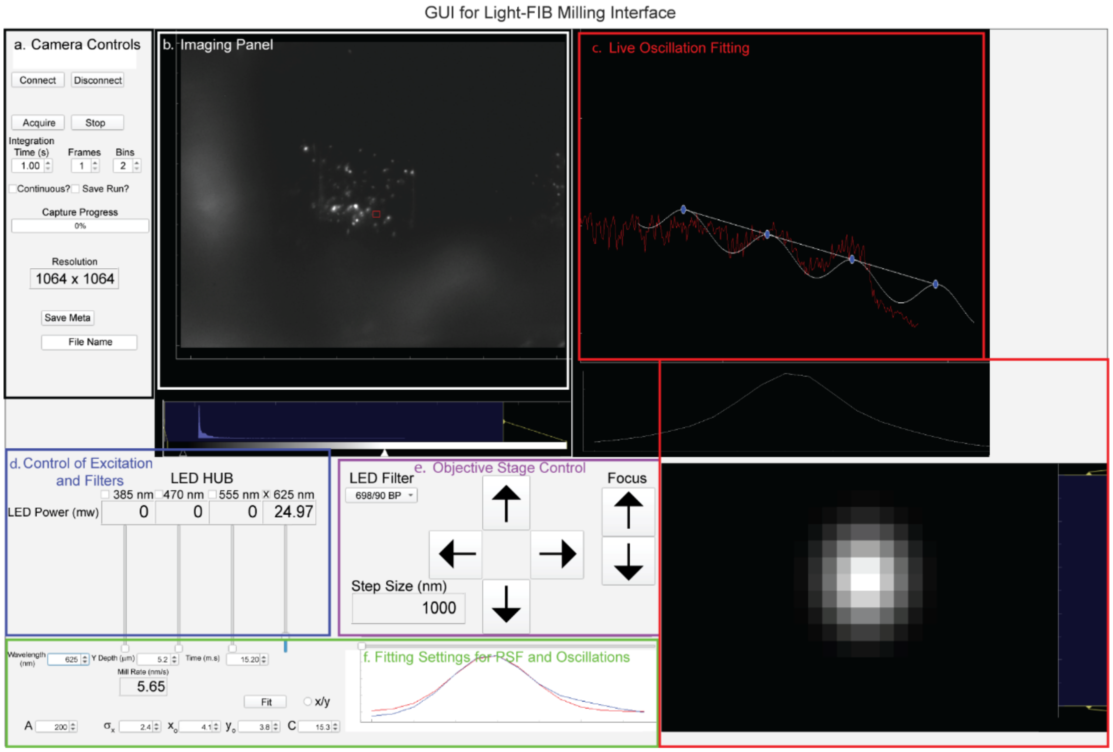


Figure S4: Custom Python-based software that provides real-time feedback on when to stop milling to capture fluorescent targets of interest. a.) Control of the camera actions and measurement parameters such as integration time. b.) Real-time optical microscopy images. Small red box identifies an ROI for PSF fitting. c.) PSF fitting result and a plot of the PSF intensity with time fit to equation 1 from the main text. d.) LED excitation controls. e.) Filter and objective stage controls. f.) Fitting parameters for the PSF and the oscillatory function in (c). When the user-defined oscillation amplitude is achieved, the user is alerted to stop milling.

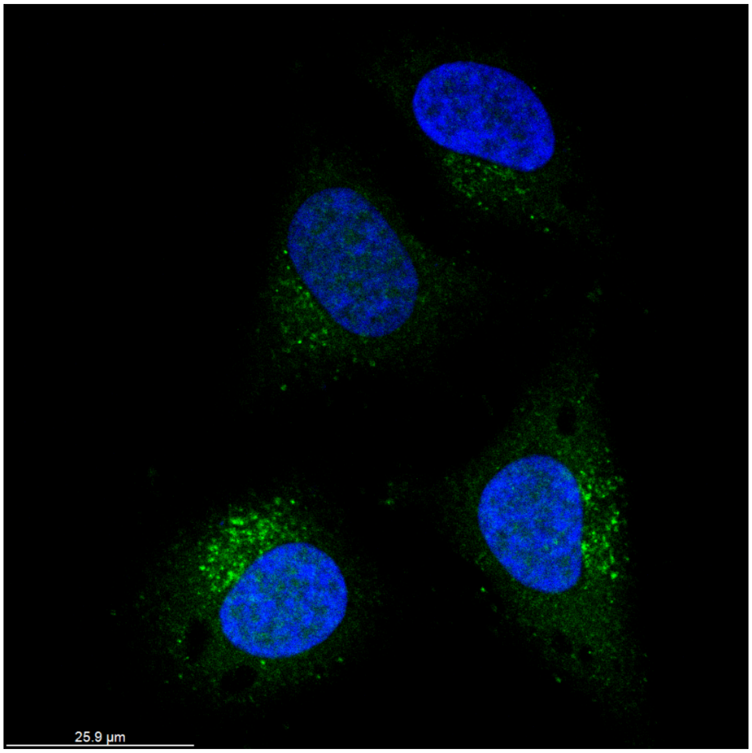


Figure S5: Confocal fluorescent microscopy section of paraformaldehyde-fixed HeLa cells infected with AAV2 non-replicating virions at an MOI of 104, shown 20 hours post infection. The virions, labeled with Alexa Fluor 488, are displayed in green, while the nuclei, stained with Hoechst, are shown in blue.
